# Chemical proteomic profiling of protein dopaminylation in colorectal cancer cells

**DOI:** 10.1101/2024.04.27.591460

**Authors:** Nan Zhang, Shuaixin Gao, Haidong Peng, Jinghua Wu, Huapeng Li, Connor Gibson, Sophia Wu, Jiangjiang Zhu, Qingfei Zheng

## Abstract

Histone dopaminylation is a newly identified epigenetic mark that plays a role in the regulation of gene transcription, where an isopeptide bond is formed between the fifth amino acid residue of H3 (*i.e.*, glutamine) and dopamine. In our previous studies, we discovered that the dynamics of this post-translational modification (including installation, removal, and replacement) were regulated by a single enzyme, transglutaminase 2 (TGM2), through reversible transamination. Recently, we developed a chemical probe to specifically label and enrich histone dopaminylation via bioorthogonal chemistry. Given this powerful tool, we found that histone H3 glutamine 5 dopaminylation (H3Q5dop) was highly enriched in colorectal tumors, which could be attributed to the high expression level of TGM2 in colon cancer cells. Due to the enzyme promiscuity of TGM2, non-histone proteins have also been identified as targets of dopaminylation on glutamine residues, however, the dopaminylated proteome in cancer cells still remains elusive. Here, we utilized our chemical probe to enrich dopaminylated proteins from colorectal cancer cells in a bioorthogonal manner and performed the chemical proteomics analysis. Therefore, 425 dopaminylated proteins were identified, many of which are involved in nucleic acid metabolism and transcription pathways. More importantly, a number of modification sites of these dopaminylated proteins were identified, attributed to the successful application of our chemical probe. Overall, these findings shed light on the significant association between cellular protein dopaminylation and cancer development, further suggesting that to block the installation of protein dopaminylation may become a promising anti-cancer strategy.

**TOC:** 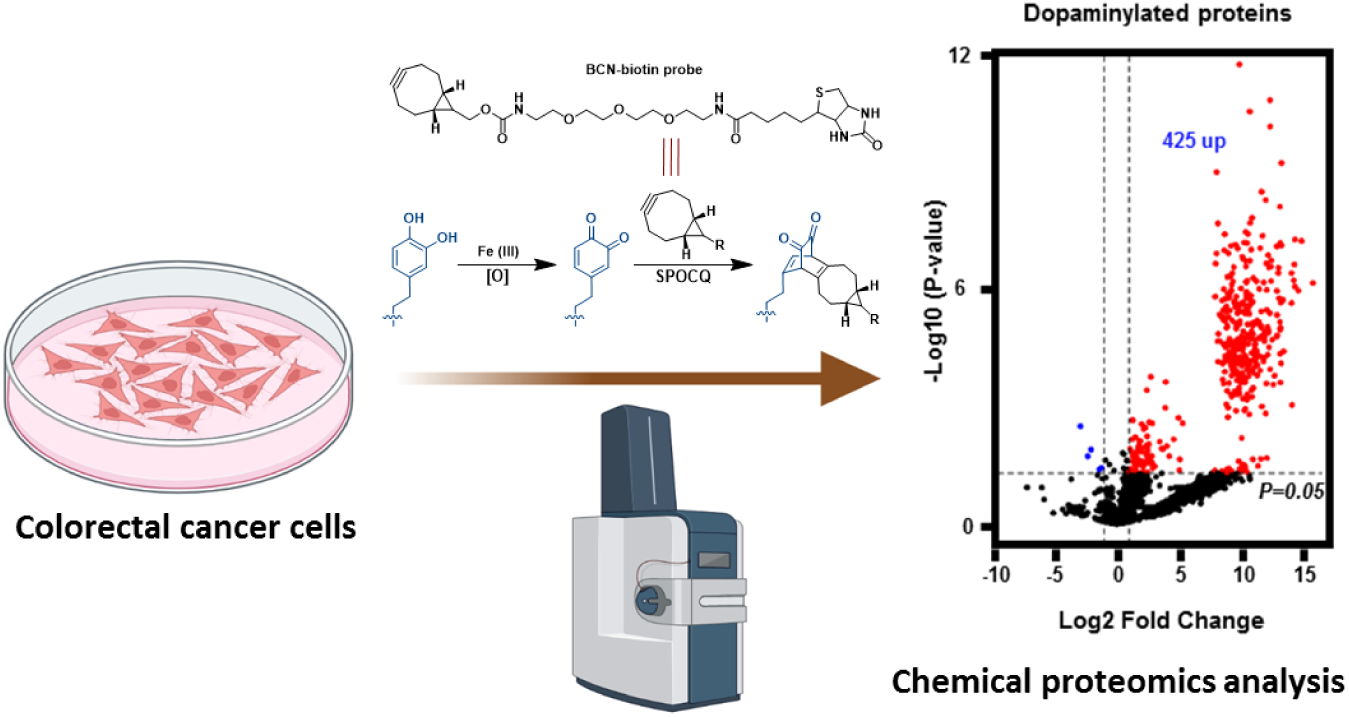

## Introduction

Protein post-translational modifications (PTMs) can significantly expand the diversity and function of cellular proteome, which play key roles in signaling transduction.^1^ Due to their fundamental functions in cell signaling,^2,3^ to target the dynamics (including the installation and removal) of protein PTMs becomes a considerable therapeutic strategy for human disease treatment.^4^ In general, the dynamics of protein PTMs are regulated by “writer” and “eraser” enzymes.^5^ In the past years, we focused on non-canonical protein PTMs that do not require writers (such as non-enzymatic glycation),^6–9^ or in which the installation and removal are regulated by a single enzyme (such as monoaminylation).^10^ Both of these two types of PTMs have been proven to serve as hallmarks in cancer cells, and thus the corresponding regulators (*i.e.*, DJ-1/PARK7,^11^ protein arginine deiminase 4,^12^ and transglutaminase 2)^10^ that are overexpressed in tumors may act as promising anti-cancer targets.^13^

In our previous studies, we found that the fifth amino acid residue (glutamine) of H3 (H3Q5) was a major modification site of transglutaminase 2 (TGM2)-mediated dopaminylation in colorectal tumors.^13^ This stable PTM is attributed to the isopeptide formation between H3Q5 and dopamine through the transamination reaction catalyzed by TGM2, which is overexpressed in colorectal cancer cells.^13^ H3Q5 dopaminylation has been shown to play a role in gene transcription regulation.^14^ Moreover, we have also uncovered that TGM2-mediated dopaminylation is a cellular microenvironment-dependent PTM, due to the reversible formation of thioester bound between TGM2 cysteine 277 (C277) and the corresponding glutamine residue of substrate protein.^10^ Given the enzyme promiscuity of TGM2,^15^ there should be other cellular proteins activated by TGM2 and undergoing dopaminylation. However, because of a lack of efficient tools (such as the pan-specific antibodies),^16^ the knowledge of dopaminylated proteins and their corresponding modification sites in cancer cells still remain poorly understood.

Currently, mass spectrometry (MS)-based proteomics is the major approach for protein PTM analysis.^1^ However, due to the low abundance of specific protein PTM and coexistence of multiple PTMs on the digested peptide fragments, the proteomic profiling of most cellular protein PTMs still stays challenging.^1^ Pan antibody-based pull-down enrichment of proteins with the specific PTM can significantly enhance the sensitivity and accuracy of proteomics analysis.^16^ Bioorthogonal chemistry-based chemical proteomics is an alternative valid method for protein PTM profiling, as the development of pan antibodies against certain PTMs cannot always be successful.^1^ Due to the lack of pan-specific antibodies and chemical tools, the proteomic profiling of protein dopaminylation in cancer cells has not been successfully performed. Recently, we have development a bicyclononyne (BCN)-containing probe that can modify and enrich dopaminylated histone proteins in a bioorthogonal manner.^13^ This BCN probe with a biotin tag enabled us to conduct the chemical proteomic profiling of protein dopaminylation in cells.

In this study, we employed the BCN-biotin probe to label and enrich the dopaminylated proteins in colorectal cancer cells via the bioorthogonal chemistry targeting dopamine residues, *i.e.*, strain-promoted oxidation-controlled cyclooctyne-1,2-quinone cycloaddition (SPOCQ; **Fig. 1A**).^13,17^ Utilizing this newly developed chemical proteomics methodology (**Fig. 1B**), we successfully identified 425 dopaminylated proteins in colon cancer cells (**Fig. 2**), which are involved in diverse signaling pathways (such as nucleic acid metabolism) of the Kyoto Encyclopedia of Genes and Genomes (KEGG) database (**Fig. 3**). Notably, a significant proportion (~72%) of these dopaminylated proteins were found to localize in the cell nucleus, many of which are related to cancer and neurodevelopment (**Fig. 4**). As the enrichment and digestion of dopaminylated proteins were of high quality, we further identified the modification sites of many key dopaminylation targets. Importantly, dopamine adducts were found on both glutamine and cysteine residues (**Fig. 5**), which are attributed to transamination and non-enzymatic covalent modification (NECM), respectively (**Fig. 5**).^7^ Overall, the chemical proteomic profiling of dopaminylated proteins in this research shed light on the ubiquitous existence of dopaminylation in cancer cells and the potential regulatory roles it plays in signaling transduction.

**Figure 1.**
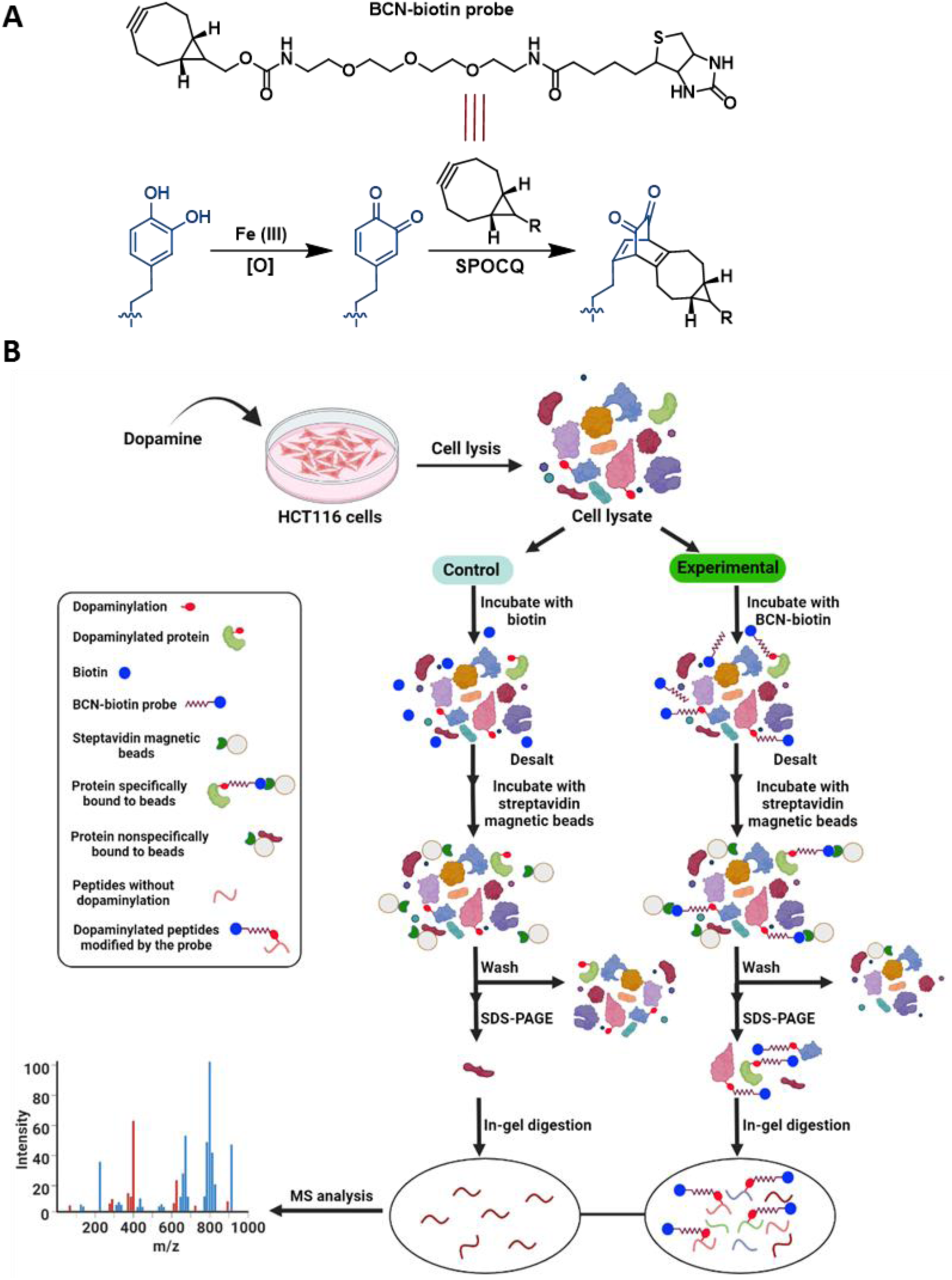
Chemical proteomic profiling of protein dopaminylation in colorectal cancer cells. (**A**) Chemical structure of the BCN-biotin probe and the bioorthogonal chemistry, strain-promoted oxidation-controlled cyclooctyne-1,2-quinone cycloaddition (SPOCQ), used in this study. (**B**) Workflow of the chemical proteomics approach developed in this study for profiling protein dopaminylation in colon cancer cells.

## Experimental section

### Colorectal cancer cell culture

HCT 116 cells (a gift from Dr. Jing J. Wang’s lab at OSU) were cultured in Dulbecco’s Modified Eagle Medium (DMEM) supplemented with 10% fetal bovine serum (FBS) and 1% sodium pyruvate under 5% CO_2_ at 37 °C. Upon reaching 80% confluence, 10 mL of medium containing 50 µL of 100 mM dopamine hydrochloride was added to achieve a final concentration of 0.5 mM. The medium was gently mixed, and cells were collected through centrifugation after 12 hours of culture. The cell pellets were washed three times using 1X Dulbecco’s Phosphate-Buffered Saline (DPBS) buffer before being further processed.

### Probe labeling and enrichment of dopaminylated proteins

The cytoplasmic and nuclear proteins of HCT 116 cells were extracted using the Cytoplasmic and Nuclear Protein Extraction Kit (BOSTER) in accordance with the provided instructions. Subsequently, 140 µL of the protein samples were separately distributed into two 1.5mL tubes, designated as the experimental and control groups, for the further probe labeling reactions. To initiate the labeling reaction, 10µL of 10X phosphate buffer (pH 8.0) was added to the reaction system. The 10X phosphate buffer (pH 8.0) was prepared by combining 1 M K_2_HPO_4_ and 1 M KH_2_PO_4_ with a volume ratio of 47:3. The protein mix was then incubated with 10 µl of K_3_[Fe(CN)_6_] (5 mM) for 5-10 minutes to oxidize the dopamine residues to dopamine quinone. Subsequently, 10 µL of BCN-biotin probe (5 mM) dissolved in DMSO was added for a one-hour reaction at room temperature, while the control sample underwent a similar reaction with 10 µL of biotin (5 mM in DMSO). Finally, the reaction was quenched by adding dopamine hydrochloride with a final concentration of 1 mM for 30 minutes. Thereafter, the samples were subjected to desalting using Zeba spin columns (Thermo Fisher Scientific) to get rid of the small molecules from the protein samples.

The streptavidin magnetic beads (Thermo Fisher Scientific) employed for enrichment underwent an initial blocking step with 1% bovine serum albumin (BSA) dissolved in 1X phosphate buffer (pH 8.0) on a rotator for 1 hour at room temperature. Subsequently, the beads were washed three times with 1X phosphate buffer for 10 minutes each. The labeled protein samples were then incubated with 40 μL streptavidin beads in 1X phosphate buffer on a rotator at 4 °C overnight to facilitate the enrichment of BCN probe-labeled proteins. Following the incubation, the beads were collected using a magnetic stand and subjected to a series of washes. Initially, the beads were washed with 1 mL of 1% NP40, 0.1% SDS, and 4 M urea, where each rotation lasted for 1 hour at room temperature. Subsequently, the beads were washed with 1X PBST (PBS containing 0.1% Tween 20 detergent) for 10 minutes to remove urea.

### SDS-PAGE analysis and silver staining

The processed streptavidin beads samples were mixed with 20 µL of 2X loading buffer (Thermo Fisher Scientific) and heated at 98℃ for 10 minutes. Subsequently, the samples were centrifuged at 16000X g for 10 minutes, and the resulting supernatant was loaded onto the SDS-gel for SDS-PAGE analysis. For visualization, silver staining was performed using the Pierce™ Silver Stain for Mass Spectrometry kit, adhering to the provided instructions.

### In-gel digestion

The protein lanes on the gel corresponding to the experimental and control samples were excised by using a surgical blade. Each gel strip was then placed into a 1.5 mL centrifuge tube and sectioned into small fragments (0.5 × 0.5 mm) using a pipette tip. The decolorizing liquid (buffer A and B in the silver staining kit) was added to immerse the gel particles until the yellow color faded, and the resulting supernatant was discarded. Following this, the gel fragments were treated with 300 μL of 50 mM NH_4_HCO_3_ for 10 minutes. After removing the liquid, 300 µL of 100% acetonitrile was added, and the tubes were placed on a shaker for 15 minutes to dehydrate the gel fragments. The supernatant was then removed, and the gel was air-dried. Thereafter, the gel fragments were incubated with a solution containing 100 mM NH_4_HCO_3_ and 10 mM tris(2-carboxyethyl)phosphine (TCEP) for 30 minutes at room temperature. The supernatant was removed, and the gel fragments were dehydrated again using 300 uL of 100% acetonitrile. Further treatment involved adding 100 μL of a solution containing 55 mM iodoacetamide in 100 mM NH_4_HCO_3_, followed by an alkylation reaction in the dark for 30 minutes. After removing the supernatant, each tube was treated with 300 μL of 100 mM NH_4_HCO_3_ at room temperature for 15 minutes on a shaker. Before enzyme lysis, another dehydration step was conducted with 300 μL of 100% acetonitrile. Each tube was then filled with 200 μL (sufficient to cover the gel particles) of a 50 mM NH_4_HCO_3_ solution containing 2 μg of trypsin (Fisher Scientific) and incubated in a temperature-controlled shaker at 37 °C for approximately 20 hours. Following enzymatic digestion, the liquid was transferred to another tube, and 150 μL of a prepared buffer (containing 5% formic acid and 50% acetonitrile) was added for peptide extraction for 20 minutes at room temperature. The tubes were shaken for 1 minute, and this extraction step was repeated three times. The final extraction was conducted using 100% acetonitrile. All the liquid was collected into a 1.5 mL centrifuge tube, followed by vacuum drying. The dried powder resulting from the previous steps was reconstituted by adding 600 µL of 0.1% formic acid aqueous solution. Subsequently, peptide desalting was carried out using peptide desalting spin columns (Pierce), following the product instructions. Finally, the desalted peptides were vacuum-dried in preparation for further LC-MS analyses.

### Mass Spectrometric Analysis

The peptide samples were dissolved in 0.1% formic acid and subjected to analysis using a hybrid trapped ion mobility spectrometry (TIMS) quadrupole time-of-flight mass spectrometer (Bruker timsTOF Pro) with a CaptiveSpray nanoelectrospray ion source coupled to a nanoflow LC system (nanoElute, Bruker). The samples were initially loaded onto a C18 column of dimensions 75 µm × 250 mm (Ionopticks). Separation was achieved at a constant flow of 300 nL min^−1^ with a linear gradient of acetonitrile, starting from 2% and reaching 22% over 94 minutes. Subsequently, the gradient was adjusted to 37% acetonitrile for 10 minutes, followed by 100% acetonitrile for 6 minutes. A final step involved maintaining 100% acetonitrile for an additional 10 minutes to ensure the analysis of all loaded peptides. Peptides were detected using a data-dependent CID method in a PASEF (Parallel Accumulation and Serial Fragmentation) mode, with 10 PASEF scans performed during each topN acquisition cycle. Single-charged precursors were excluded based on their positions in the m/z-ion mobility plane. Precursors reaching the target value of 20000 a.u. were dynamically excluded for 0.4 minutes. For m/z values less than 700, the quadrupole isolation width was set to 2 Th, while for m/z values greater than 700, the quadrupole isolation width was set to 3 Th. The accumulation and ramp times were both set to 100 ms, and mass spectra were recorded in the positive electrospray mode within the m/z range of 100 to 1700. Ion mobility was scanned from 0.6 to 1.6 V s/cm². During the scanning mode, collision energy underwent a linear ramp as a function of mobility, ranging from 59 eV at 1/K₀ = 1.6 V s/cm²to 20 eV at 1/K₀ = 0.6 V s/cm².

### Data processing and statistical analysis

Fragpipe software (v21.1) was utilized for processing the MS and MS/MS raw data. The MS/MS spectra were aligned with the human UNIPROT database, which comprises 20,421 human reviewed entries. The applied search criteria included strict trypsin digestion and the incorporation of variable modifications such as glutamine and cysteine dopaminylation modified by the BCN-biotin probe (+728.3455 Da; +136.0524 Da; +134.0368 Da; +744.3642 Da; +740.3329 Da; +152.0712 Da; +150.0555 Da; +146.0242 Da), as well as cysteine carboxyamidomethylation (+57.0214 Da). The analysis allowed for up to three missed cleavages, and other search parameters were set to default values. For statistical analysis, Omicsbean software was employed. A gene filter was applied to calculate the fold-change values of protein expression. The t-test analysis was utilized to filter differentially expressed proteins, with a fold change threshold set at greater than 2-fold or less than 0.5-fold, and a significance level (P-value) below 0.05.

## Results and discussion

### Probe labeling and enrichment of dopaminylated proteins from HCT116 cells

The BCN-biotin probe (**Fig. 1A**) was synthesized (**Supplementary Information**) and characterized, based on the protocols reported in our previous study.^13^ HCT116 cells were cultured and utilized as a model of colorectal cancer to identify dopaminylated proteins. As TGM2 is overexpressed in colon cancer cells,^13^ the experimental and control groups were treated with the same amount of dopamine to induce dopaminylation on the target proteins. The cellular proteins were extracted and then treated with Fe (III) to oxidize the dopamine residues of modified proteins to dopamine quinone, which can be modified by the BCN-biotin probe via a bioorthogonal [4 + 2] cycloaddition.^13,17^ In the control groups, the same amount of biotin was added to the cell lysates instead of the probe (**Fig. 1B**). The streptavidin magnetic beads were utilized to enrich the probe-modified proteins. Further SDS analysis and silver staining indicated that dopaminylated proteins were obviously and robustly enriched by using the BCN-biotin probe-based pull-down in the experimental groups (**Fig. S1**).

### Profiling of dopaminylated proteome in HCT116 cells

The collected raw MS data first underwent standardization for subsequent analyses (**Fig. S2** and **Supplementary Data 1-3**). Distinct stratification among different groups was observed through Principal Components Analysis (PCA) (**Fig. S3**). The comparison analysis further indicated that there were 425 proteins significantly enriched in the BCN-biotin probe-treated group (**Fig. 2A**). Notably, these experiments were biologically repeated in triplicate and the data of enriched proteins were consistent (**Fig. 2B**). Further signaling pathway analysis using the Kyoto Encyclopedia of Genes and Genomes (KEGG) database indicated that the enriched proteins with dopaminylation were involved in more than ten pathophysiologically important pathways, with substantial impacts on the spliceosome, RNA transport, endocrine, choline metabolism, protein processing in the endoplasmic reticulum (ER), *etc* (**Fig. 2C**). Based on the Biological Process (BP)-KEGG analysis, it showed that the enriched KEGG pathways were predominantly associated with three biological processes: nucleic acid-templated transcription, RNA splicing, and nervous system development (**Figs. 2D** and **S4**). Specifically, 305 (~72%) of the enriched proteins are localized in the nucleus and 405 (~96%) of them are associated with the molecular function related to biomacromolecule binding (**Figs. 2E** and **2F**). These significantly enriched nuclear binding proteins (BPs) play crucial roles in various cellular functions, including gene transcription, protein translation, DNA repair, RNA splicing, apoptosis, stress responses, *etc*. The results of these analyses suggested that dopamine might play indispensable roles in modulating the key biological processes within nucleus by modifying the amino acid residues of nuclear BPs.

**Figure 2.**
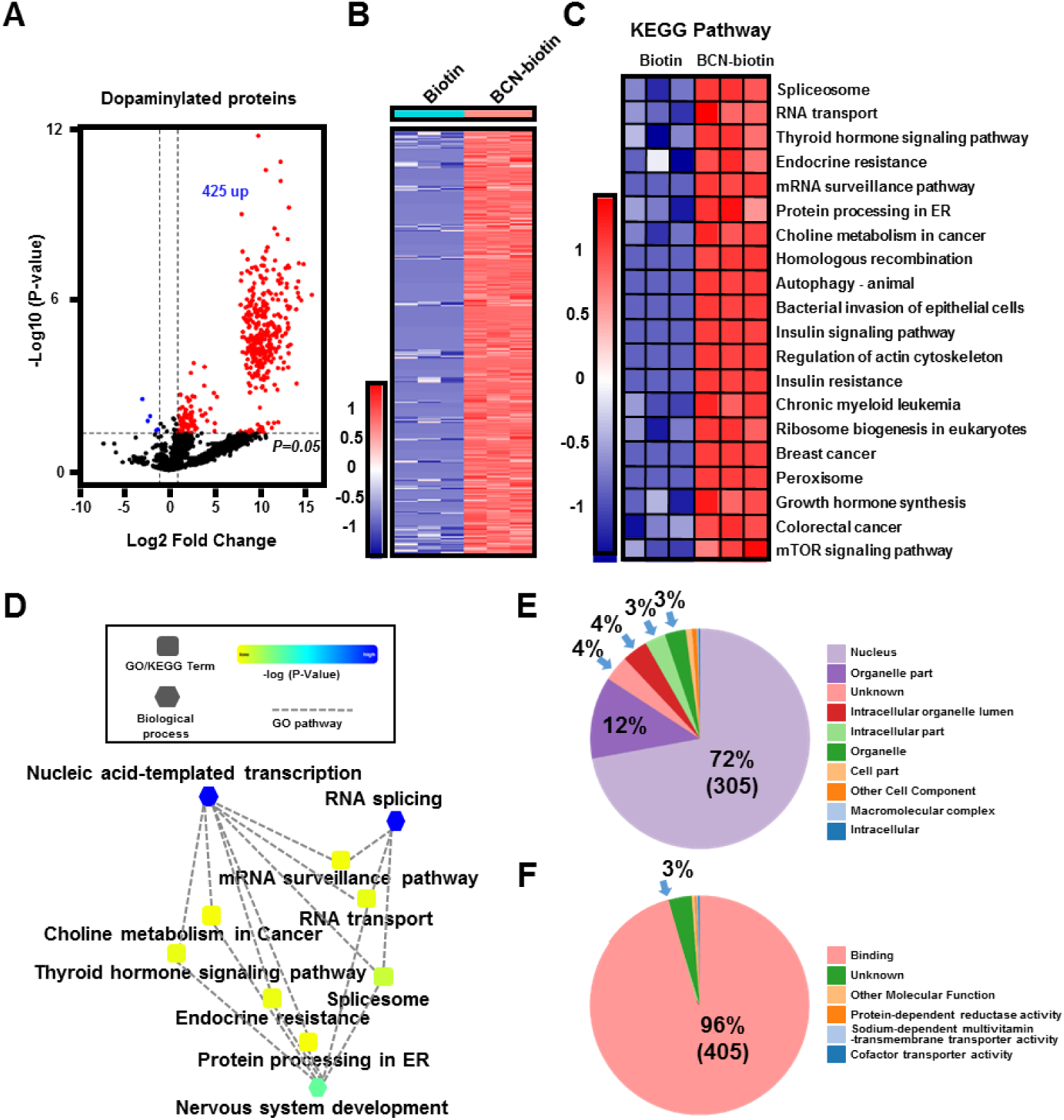
Comprehensive profiling of the dopaminylation proteome in HCT116 cells. (**A**) T-test comparison between the BCN-biotin probe-enriched and biotin-treated proteins. The analysis encompassed the data from three independent experiments. (**B**) Heatmap representation of label-free quantitation (LFQ) intensity data of the 425 dopaminylated proteins enriched by using the BCN-biotin probe. The heatmap color (from red to blue) represents the fold change of protein level from increasing to decreasing. (**C**) Heatmap representation and functional annotation of the clusters derived from KEGG (Kyoto Encyclopedia of Genes and Genomes) pathways. (**D**) Network showing the link between biological processes and KEGG pathways of the identified proteins. (**E**) Cell component of the identified proteins possessing dopaminylation. (**F**) Molecular functions of the identified proteins possessing dopaminylation.

### Interaction network analysis of enriched proteins containing dopaminylation

Notably, 298 (~98%) of the 305 enriched proteins that contain dopaminylation and localize to the cell nucleus belong to the group of proteins with binding functions (**Fig. 3A**). Moreover, 208 of these proteins are associated with nucleic acid metabolic processes (**Fig. 3A**), encompassing primarily ten biological processes: mRNA processing, protein modification by small protein conjugation, regulation of mRNA metabolic process, peptidyl-lysine modification, protein export from nucleus, RNA splicing, nucleic acid-template transcription, RNA 3’-end processing, and protein phosphorylation regulation (**Fig. S4**). Further clustering analysis indicated that two of these biological processes related to the nucleic acid metabolism, *i.e.*, RNA splicing^18^ and nucleic acid-template transcription,^19^ are particularly emphasized, with a significant protein cluster comprising 176 (~85%) of the 208 nucleic acid metabolic process-related proteins that contain dopaminylation (**Figs. 3A, 3B**, and **S4**). The molecular functions of these 176 RNA splicing and nucleic acid-template transcription-related proteins could be further broken down into nucleic acid binding, transcription factor binding, transcription cofactor binding, sequence-specific DNA binding, identical protein binding, protein domain-specific binding, RNA polymerase II transcription factor binding, enzyme binding, modification-dependent protein binding, histone binding, *etc* (**Fig. 3C**). As anticipated, this cluster of proteins includes various transcription factors, such as zinc finger proteins (ZNF16, ZNF217, ZNF512B, ZNF787, ZNF579, ZNF668, ZNF629),^20^ mothers against decapentaplegic homolog 4 (SMAD4), BTB domain-containing protein 7A (ZBTB7A), and transcriptional repressor protein YY1 (YY1), all of which exhibit molecular functions associated with protein binding and nucleic acid binding (**Fig. 3D**).^21^ Overall, all of these results indicate that protein dopaminylation plays an essential role in RNA splicing and nucleic acid-template transcription processes within the cell nucleus.

**Figure 3.**
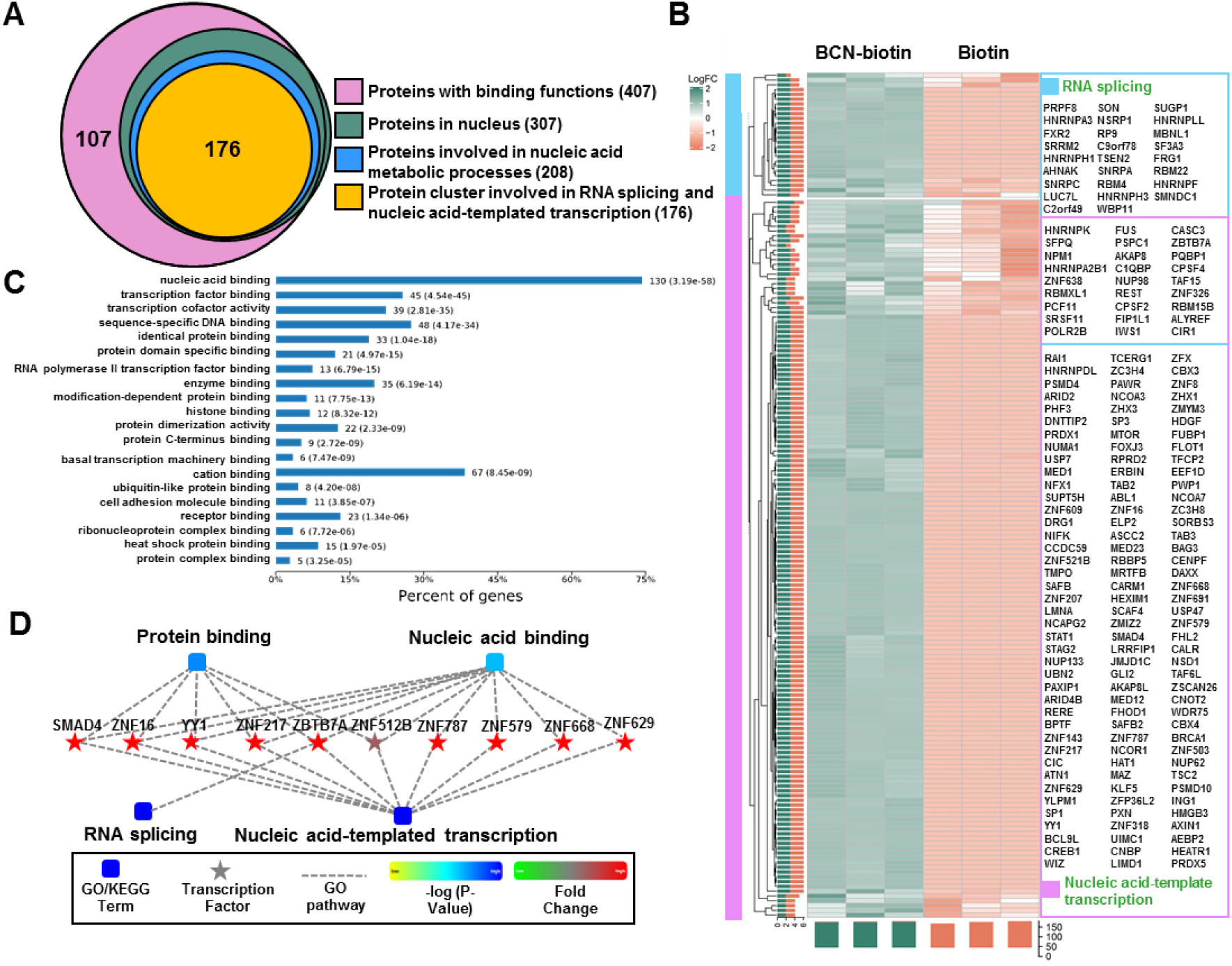
Function analysis of the dopaminylation proteome of HCT116 cells. (**A**) Venn diagram showing a prominent cluster containing 176 dopaminylated proteins that are associated with RNA splicing and nucleic acid-templated transcription. (**B**) Heatmap indicating the 176 dopaminylated proteins that are involved in RNA splicing and nucleic acid-templated transcription pathways. (**C**) Molecular functions of the 176 dopaminylated proteins in the prominent cluster. (**D**) Network diagram depicting the key transcription factors identified in the prominent cluster of dopaminylated proteins. Color bar from green to red represents the fold change of protein levels from increasing to decreasing. The significance of pathways represented by −log(p value) (Fisher’s exact test) was shown by color scales with dark blue as the most significant.

To uncover the pathophysiological relevance of protein dopaminylation, we conducted further interaction network analysis on the cluster of aforementioned 176 dopaminylated proteins and observed that a number of them were associated with carcinogenesis and cancer development (**Fig. 4A**). Specifically, the interaction diagram illustrated that these RNA splicing and nucleic acid-template transcription-related proteins possessing dopaminylation involved many key regulators in cancer development, such as transcription factors Sp1 (SP1),^22^ serine/threonine-protein kinase mTOR (MTOR),^23^ tyrosine-protein kinase ABL1 (ABL1),^24^ heterogeneous nuclear ribonucleoproteins (hnRNPs),^25^ *etc*, which are druggable targets for cancer therapies. Additionally, among these dopaminylated proteins, the transcriptional repressor protein YY1 and Aly/REF export factor (ALYREF) have been identified as potential tumor biomarkers for both diagnosis and prognosis.^26^ In the cancer-related cluster (**Fig. 4A**), there are also numerous proteins playing key roles in carcinogenesis,^27^ which include the splicing factor polyglutamine-binding protein 1 (PQBP1),^28^ small nuclear ribonucleoprotein polypeptide C (SNRPC),^29^ small nuclear ribonucleoprotein polypeptide A (SNRPA),^29^ RNA-binding protein FUS (FUS),^30^ histone-arginine methyltransferase CARM1 (CARM1),^31^ splicing factor proline- and glutamine-rich (SFPQ) protein,^28^ nuclear receptor coactivator 3 (NCOA3),^32^ nuclear pore glycoprotein p62 (NUP62),^33^ *etc*.

**Figure 4.**
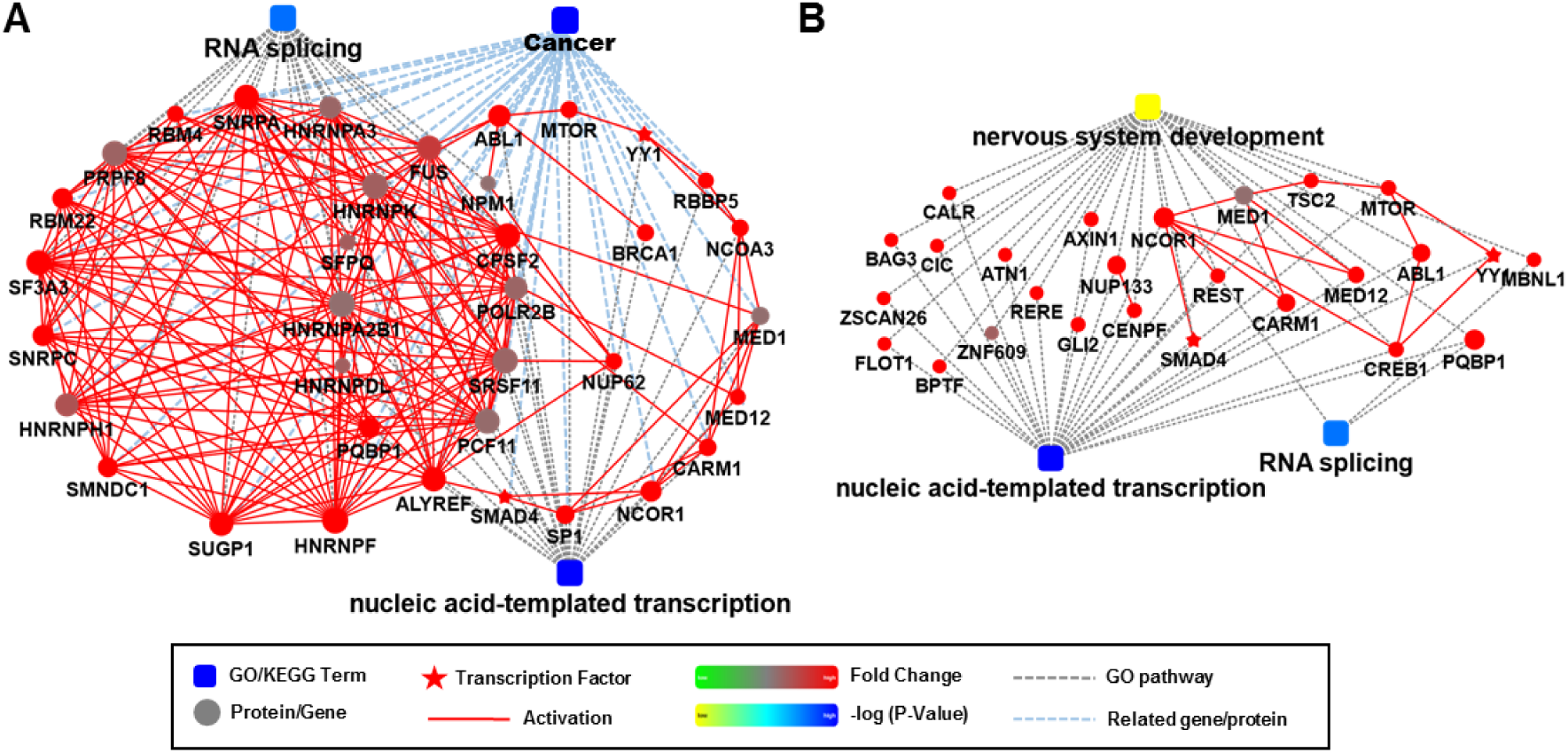
Interaction diagram illustrating two highly enriched groups, (**A**) cancer and (**B**) nervous system development, within the prominent dopaminylated cluster. Color bar from green to red represents the fold change of protein levels from increasing to decreasing. The significance of the pathways represented by −log(p value) (Fisher’s exact test) was shown by color scales with dark blue as most significant.

Moreover, within the cluster of identified 176 dopaminylated proteins that are associated with RNA splicing and nucleic acid-template transcription, a subset was discovered to be linked to nervous system development (**Fig. 4B**), which further highlighted the essential role of dopamine in neurodevelopment as a key neurotransmitter. For example, these neurodevelopment-related proteins included zinc finger protein 609 (ZNF609),^34^ tuberous sclerosis complex 2 (TSC2),^35^ RE1-silencing transcription factor (REST),^36^ polyglutamine-binding protein 1 (PQBP1),^37^ centromere protein F (CENPF),^38^ nucleosome-remodeling factor subunit BPTF (BPTF),^39^ muscleblind-like protein 1 (MBNL1),^40^ *etc*. Additionally, there were other important regulators in neurobiology identified, including axin-1 (AXIN1) that participates in the blood-brain barrier (BBB) protection and improvement of neurological functions during ischemic stroke.^41^ In summary, the interaction network analysis indicated that there were a number of cancer and neurodevelopment-associated regulators identified to undergo dopaminylation. Further studies are needed to shed light on the impacts (such as activation or suppression) of dopaminylation on the activities of these pathophysiologically important proteins.

### Identification of dopaminylation sites

Previous studies have shown that protein dopaminylation can occur in two different manners: 1) TGM2-mediated dopaminylation on glutamine (Q) residues;^10,14^ 2) dopamine quinone-induced non-enzymatic dopaminylation on cysteine (C) residues (**Fig. 5A**).^42^ Notably, these two types of dopaminylation are both able to be bioorthogonally modified by our BCN-biotin probe *via* SPOCQ (**Fig. 1A**). As the Diels-Alder reaction between BCN and dopamine quinone is reversible under the ionization conditions of mass spectrometer, there are eight possible molecular weight (MW) shifts in the modified peptide fragments (**Fig. 5**) and the two types of dopaminylation can be distinguished based on the different mass shifts. The dopaminylation sites (Q or C) of many key regulators, such as transforming acidic coiled-coil-containing protein 2 (TACC2),^43^ scaffold attachment factor B (SAFB),^44^ peroxiredoxin-4 (PRDX4),^45^ zyxin (ZYX),^46^ *etc*, were identified based on the high-quality mass spectra (**Fig. 5** and **Table S1**). Importantly, the observation of distinct modification residues and mass shifts on the same peptide fragment (SAFB in **Fig. 5B**) further confirmed the occurrence of dopaminylation on this specific amino acid site. Overall, the dopaminylation on these sites may significantly influence the activities of these pathophysiologically important proteins *via* the allosteric effect, steric effect, electronic effect, π-π stacking, or directly blocking the key catalytic sites (such as cysteine).^1,13^ For example, PRDX4 is a cysteine-dependent antioxidant enzyme, of which the key cysteine dopaminylation (**Fig. 5B**) may destroy the protein disulfide bond structure and abolish the enzymatic activity.^47^ Further studies are required to uncover the pathophysiological implications of the dopaminylation on these sites.

**Figure 5.**
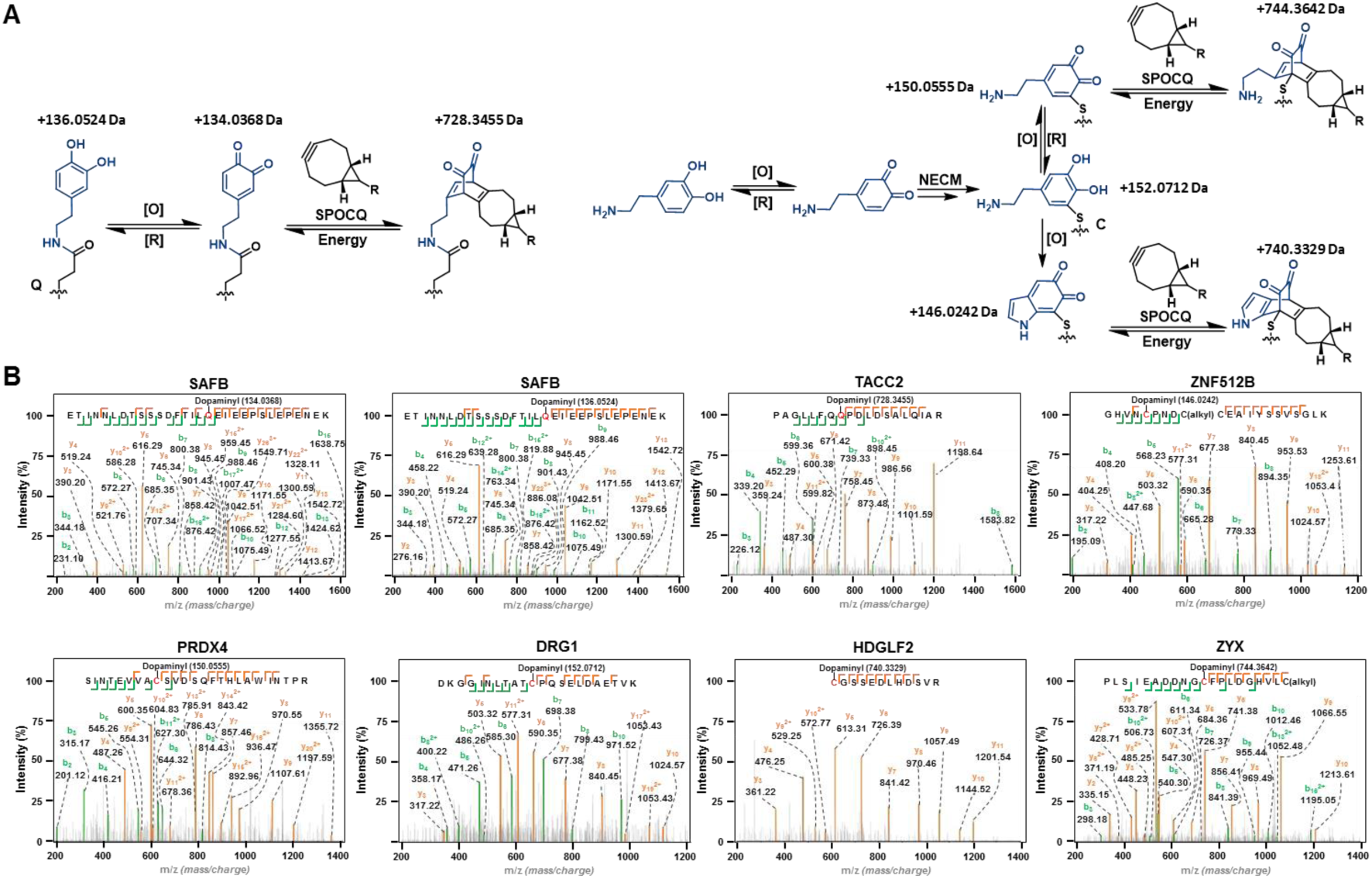
Modification site analysis of the enriched proteins containing dopaminylation in HCT116 cells. (**A**) Fragment structures of the dopaminylated proteins (on Q and C) and the corresponding mass shifts. (**B**) Examples of the identified modification sites (on Q and C) and the corresponding mass shifts.

## Conclusion

Dopamine is a metabolite converted from the essential amino acid, phenylalanine,^48^ which plays significant regulatory roles in cellular signaling transduction as a well-known neurotransmitter.^49^ The discovery of protein dopaminylation added a novel mode of action (MOA) to dopamine-mediated signaling pathways. One well-established example is that histone H3 dopaminylation (H3Q5dop) was proven to be implicated in cocaine-induced transcriptional plasticity, heroin-induced transcriptional and behavioral plasticity, as well as drug-induced transcriptional and behavioral changes.^14,50^ In our previous studies, we have uncovered that the dynamics of histone H3 dopaminylation is regulated by the single enzyme, TGM2.^10^ Due to the enzyme promiscuity of TGM2, there are diverse non-histone proteins undergoing dopaminylation on glutamine residues. Moreover, cysteine residues can also be modified by dopamine quinone via non-enzymatic dopaminylation.^42^ Even though protein dopaminylation (such as the newly identified epigenetic mark, H3Q5dop) has been shown to play essential roles in signaling pathways (*e.g.*, gene transcription),^14^ the cellular dopaminylated proteome is still poorly understood.

Recently, we designed and developed a BCN-based chemical probe to label and enrich dopaminylated proteins in a biorthogonal fashion, which could be employed as a pan-specific antibody to study dopaminylation.^13^ Utilizing this chemical biology tool, we found that histone dopaminylation was accumulated in tumor cells that overexpress TGM2 (*e.g.*, colon and breast cancer cells).^13^ In this research, we applied this BCN-biotin probe to modify the dopaminylated proteome in colorectal cancer (HCT116) cells via the biorthogonal reaction, SPOCQ. Therefore, 425 dopaminylated proteins were significantly enriched and identified from the experimental group. Further analysis indicated that these proteins were involved in diverse fundamental pathways, most of which were found to be in the nucleus and associated with nucleic acid metabolic processes. Thereafter, our bioinformatic analyses revealed that these dopaminylated proteins were primarily related to transcriptional regulation, including RNA splicing and nucleic acid-templated transcription. Furthermore, the interaction network analyses indicated that most of the identified proteins possessing dopaminylation were closely related to cancer and neurodevelopment, highlighting the significant pathophysiological relevance of this novel PTM. Finally, utilizing the chemical proteomics approach developed in this study, we successfully identified a number of dopaminylation sites (including Q and C residues) from various pathophysiologically important proteins. In summary, we applied a novel BCN-biotin probe to perform the chemical proteomic profiling of protein dopaminylation in colon cancer cells and successfully identified over 400 dopaminylated proteins that are associated with cancer and neurodevelopment. Overall, the dataset generated by using our new methodology highlighted the importance of dopaminylation in cellular processes and provided new insights for future disease therapeutics, especially cancer and neurological disorders.

## Acknowledgements

This research work was financially supported by the NIH (R35 GM150676) and OSUCCC startup funds for Q.Z. The Zhu lab is supported by the NIH grant (R35 GM133510) for J.Z. We acknowledge Dr. Sophie Harvey and the Campus Chemical Instrument Center (CCIC) Mass Spectrometry and Proteomics Facility at OSU for their assistance in mass spectroscopy. The timsTOF Pro instrument was supported by the NIH Grant (S10 OD026945).

