## Supplemental Information for "Chemical proteomic profiling of protein dopaminylation in colorectal cancer cells"

### MATERIALS AND METHODS

#### General methods (equipment, reagents, chemicals)

UV spectrometry was performed on a NanoDrop 2000c (Thermo Scientific). Biochemicals and media were purchased from Fisher Scientific or Sigma-Aldrich Corporation unless otherwise stated. Centrifugal filtration units were purchased from Millipore, and MINI dialysis units purchased from Pierce. Gels were imaged on an Odyssey CLx Imaging System (Li-Cor). All experiments in this research were performed at least 3X. For the synthesis of probe molecules used in this study, all commercial chemicals were purchased from Sigma Aldrich, TCI chemicals, AK Scientific, Fischer Scientific, Broadpharm and used without further purification. The reagents and solvents were handled following the safety processes required as instructed by the manufacturer. Organic and aqueous waste was disposed of following standard safety protocols. Solvents for workup were purchased from Fisher Chemical, and anhydrous solvents were purchased from Sigma Millipore in a sealed bottle and degassed by passing $N_2$ before each use. Reaction progress was monitored by using normal phase TLC silica gel 60 $F_{254}$ plates by Sigma Aldrich (aluminum backed 20 X 20 cm). Developed plates were analyzed by visualizing under a UV-light and/or staining with Phosphomolybdic Acid (PMA) Stain (100 mL absolute ethanol and 10 g PMA). Isolation and purification of the crude reaction materials were performed using silica gel ($SiO_2$) by Acros Organic (0.030-0.200 mm, 60 $A^0$). Organic solvents were removed under vacuum using a Heidolph Rotavapor equipped with a dry ice condenser.

#### Synthesis of the BCN-biotin probe

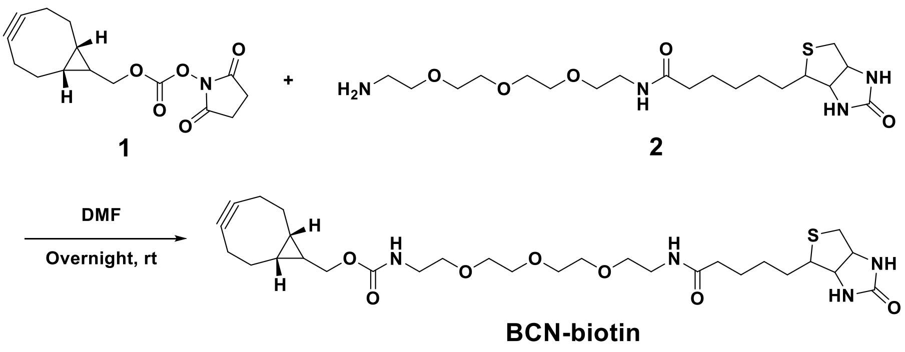


The BCN-biotin probe was synthesized based on the protocols reported in our previous study.[1] The starting material, compound **1** (100 mg, 0.3432 mmol, 1 eq), was dissolved in DMF (3 mL). Thereafter, biotin-PEG$_3$-amine, **2** (172.44 mg, 0.41 mmol, 1.2 eq), was added into the solution. The reaction mixture was washed with water (3 X 10 mL) and extracted with DCM (3 X 10 mL) after stirring overnight at room temperature. The organic layers were dried over $Na_2SO_4$ and then concentrated under reduced pressure. Preparative RP-HPLC was used to purify the BCN-biotin probe for the NMR/HR-LCMS analysis and described applications in this study.

#### Statistics and reproducibility

All the mass spectrometry analyses were repeated independently at least three times with similar results. Significance was determined at $p<0.05$. All data are represented as mean $\pm$SEM. Statistical analyses were performed in GraphPad Prism 9.

#### SUPPLEMENTARY TABLES

| Peptide Sequence | Verified sites | Gene Name |
|---|---|---|
| SSSHLPANSQDVQGYITNQSPESIVEEAQGK | 19Q(134.0368) | ANKRD17 |
| QGCPFQPWDGLEEHSQALSGR | 1Q(134.0368) | APOBEC3B |
| DNLAQQSFNMEQANYTIQSLK | 12Q(134.0368) | CHMP5 |
| DNLAQQSFNMEQANYTIQSLK | 12Q(136.0524) | CHMP5 |
| IDQIEDLQDQLEDMMEDANEIQEALSR | 8Q(134.0368) | CHMP5 |
| PTQGASSASEPQEAPPKPAEDK | 12Q(728.3455) | CLPTM1 |
| QEIIASVDHIK | 1Q(134.0368) | CPSF4 |
| FELPMGTTEQPPLPQQTQPPAK | 10Q(136.0524) | CPSF4 |
| PLAEGPQVTGPIEVPAAR | 7Q(134.0368) | CRIP2 |
| PLAEGPQVTGPIEVPAAR | 7Q(136.0524) | CRIP2 |
| DKGGINLTATCPQSELDAETVK | 13Q(134.0368) | DRG1 |
| TVEHGFPNQPSALAFDPELR | 9Q(728.3455) | LLGL1 |
| PCNSQPSELSSETSGIARPEEGRPVVSGTGNDITTPPNK | 5Q(134.0368) | MAP4 |
| DEAAGGAAAAAAEAGAASGEQAAAPGEEAAAGEEGAAGGDPQEA | 21Q(728.3455) | MARCKS |
| SILTSTTTVEHAPIWRPGTEQSSGSSGGGGGSSSR | 21Q(134.0368) | NCOR2 |
| SILTSTTTVEHAPIWRPGTEQSSGSSGGGGGSSSR | 21Q(136.0524) | NCOR2 |
| NSFYMGTCQDEPEQLDDWNR | 9Q(134.0368) | NUMA1 |
| GHTDSVQDISFDHSGK | 7Q(136.0524) | PAFAH1B1 |
| SWAQASVTHGAHGDGGR | 4Q(134.0368) | PRRC2A |
| GAAAADLLSSSPESQHGGTQSPGGGQPLLQPTK | 15Q(134.0368) | PTPN23 |
| ETINNLDTSSSDFTILQEIEEPSLEPENEK | 17Q(134.0368) | SAFB |
| ETINNLDTSSSDFTILQEIEEPSLEPENEK | 17Q(136.0524) | SAFB |
| SEPVKEESSELEQPFAQDTSSVGPDR | 17Q(134.0368) | SAFB |
| IKGEQEKELSK | 5Q(134.0368) | SLK |
| VDEDSAEDTQSNDGKEVVEVGQK | 10Q(134.0368) | SLK |
| LHQLSGSDQLESTAHSR | 9Q(134.0368) | SPAG9 |
| AGMSSNQSISSPVLDAVPR | 7Q(136.0524) | SRRM2 |
| PAGLLFQQPDLDSALQIAR | 7Q(136.0524),8Q(136.0524 | TACC2 |
| PAGLLFQQPDLDSALQIAR | 8Q(728.3455) | TACC2 |
| TDADSESDNSDNNTIFVQGLGEGVSTDQVGEFFK | 18Q(134.0368) | TAF15 |
| QDVESCYFAAQTM[15.9949]K | 11Q(136.0524) | TNPO3 |
| LGHPEALSAGTGSPQPPSFTYAQQR | 15Q(728.3455) | ZYX |
| PLSIEADDNGCFPLDGHVLCR | 11C(744.3642) | ZYX |
| SQSPAASDCSSSSSSASLPSSGR | 9C(146.0242) | BAG3 |
| SQSPAASDCSSSSSSASLPSSGR | 9C(740.3329) | BAG3 |
| YKDLEQQDCEIAQEIQEK | 9C(146.0242) | CCDC50 |
| HELQANCYEEVKDR | 7C(152.0712) | CFL1 |
| CGESGHLAKDCDLQEDACYNCGR | 1C(152.0712) | CNBP |
| DKGGINLTATCPQSELDAETVK | 11C(152.0712) | DRG1 |
| CGSSEDLHDSVR | 1C(740.3329) | HDGFL2 |
| DLNYCFSGMSDHR | 5C(152.0712) | HNRNPH1 |
| DLNYCFSGMSDHR | 5C(740.3329) | HNRNPH1 |
| NVCLPPEMEVALTEDQVPALK | 3C(146.0242) | MAP4 |
| CSLPAEEDSVLEK | 1C(740.3329) | MAP4 |
| NTHCSSLPHYQK | 4C(146.0242) | MATR3 |
| GYPHLCSICDLPVHSNK | 9C(152.0712) | MATR3 |
| HGEVCPAGWKPGSETIIPDPAGK | 5C(146.0242) | PRDX4 |
| SINTEVVACSVDSQFTHLAWINTPR | 9C(146.0242) | PRDX4 |
| SINTEVVACSVDSQFTHLAWINTPR | 9C(150.0555) | PRDX4 |
| ALNVEPDGTGLTCSLAPNIISQL | 13C(146.0242) | PRDX5 |
| SNELGDVGVHCVLQGLQTPSCK | 11C(150.0555) | RNH1 |
| PGHLQEGFGCVVTNR | 10C(744.3642) | SERBP1 |
| PGHLQEGFGCVVTNR | 10C(740.3329) | SERBP1 |
| ENFALECSPAFQVSDDEHEK | 7C(150.0555) | SGO2 |
| ENFALECSPAFQVSDDEHEK | 7C(152.0712) | SGO2 |
| ADMLCNSQSNDILQHQGSNCGGTSNK | 5C(146.0242) | TXLNG |
| TSHSSTEACCELCGLYFENR | 9C(146.0242) | WIZ |
| GHVNCPNDCCEAIYSSVSGLK | 5C(146.0242) | ZNF512B |
| GHVNCPNDCCEAIYSSVSGLK | 5C(150.0555) | ZNF512B |
| AYHPHCFTCVVCAR | 9C(146.0242) | ZYX |

Dopaminylated glutamine and cysteine residues are marked with red.

**Table S1.** Verified dopaminylation sites and the corresponding peptide sequences.

#### SUPPLEMENTARY FIGURES AND LEGENDS

**Figure S1.** Distribution map of the data in each sample. Control groups (11, 13 and 15) and experimental groups (12, 14 and 16) before and after standardization.

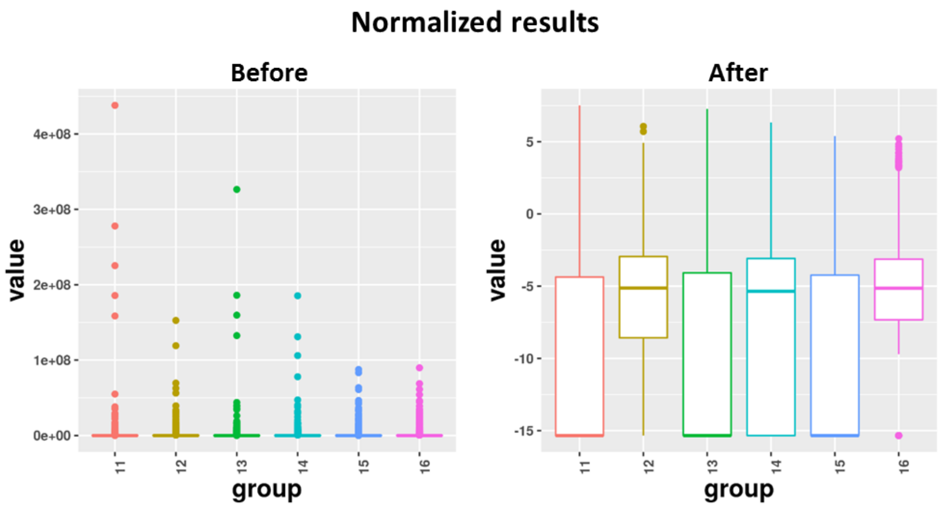

**Figure S2.** Principal Components Analysis (PCA) showed the group difference after standardization. A clear stratification shows that the two groups can be well distinguished.

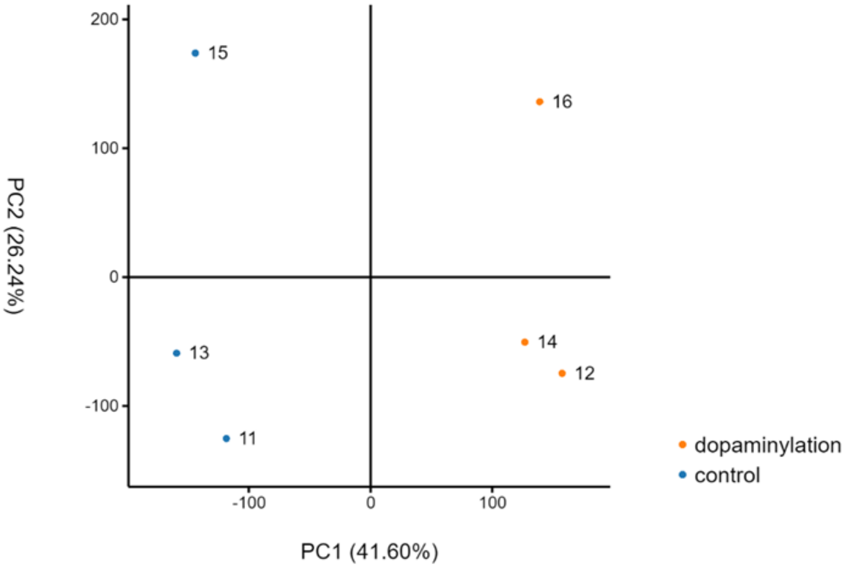

**Figure S3.** Complex heatmap of the cluster containing 176 dopaminylated proteins that are involved in the RNA splicing and nucleic acid-templated transcription pathways.

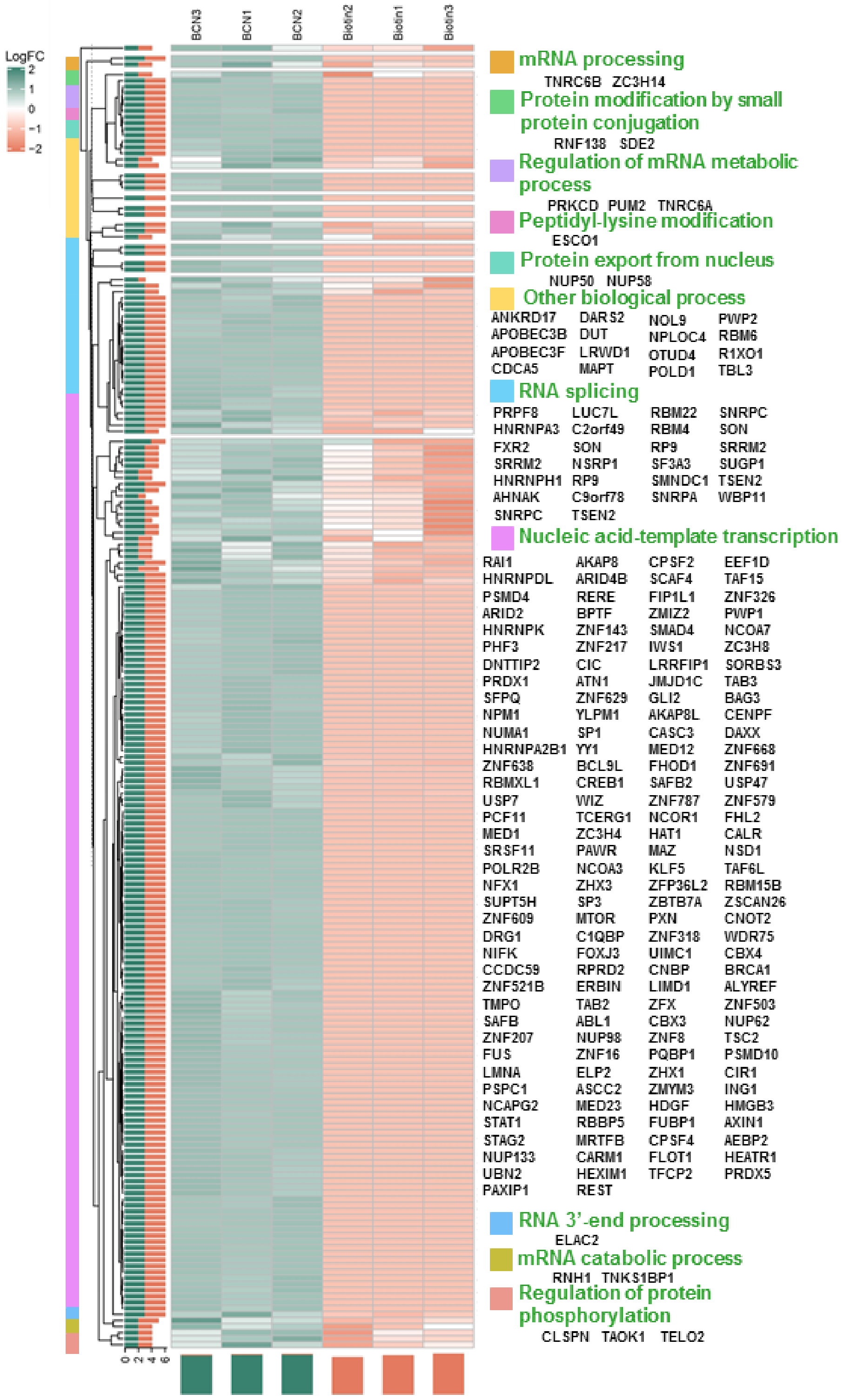


**Biological process of proteins in nucleic acid metabolic process**
